# *In vivo* imaging of the kinetics of microglial self-renewal and maturation in the adult visual cortex

**DOI:** 10.1101/2020.03.05.977553

**Authors:** Monique S. Mendes, Jason Atlas, Zachary Brehm, Antonio Ladron-de-Guevara, Matthew N. McCall, Ania K. Majewska

## Abstract

Microglia are the resident immune cells in the brain with the capacity to autonomously self-renew. Under basal conditions, microglial self-renewal appears to be slow and stochastic, although microglia have the ability to proliferate very rapidly following depletion or in response to injury. Because microglial self-renewal has largely been studied using static tools, the mechanisms and kinetics by which microglia renew and acquire mature characteristics in the adult brain are not well understood. Using chronic *in vivo* two-photon imaging in awake mice and PLX5622 (Colony stimulating factor 1 receptor (CSF1R) inhibitor) to deplete microglia, we set out to understand the dynamic self-organization and maturation of microglia following depletion in the visual cortex. We confirm that under basal conditions, cortical microglia show limited turnover and migration. Following depletion, however, microglial repopulation is remarkably rapid and is sustained by the dynamic division of the remaining microglia in a manner that is largely independent of signaling through the P2Y12 receptor. Mathematical modeling of microglial division demonstrates that the observed division rates can account for the rapid repopulation observed *in vivo*. Additionally, newly-born microglia resemble mature microglia, in terms of their morphology, dynamics and ability to respond to injury, within days of repopulation. Our work suggests that microglia rapidly self-renew locally, without the involvement of a special progenitor cell, and that newly born microglia do not recapitulate a slow developmental maturation but instead quickly take on mature roles in the nervous system.

**Graphical Abstract:** (a) Microglial dynamics during control condition. Cartoon depiction of the heterogenous microglia in the visual cortex equally spaced. (b) During the early stages of repopulation, microglia are irregularly spaced and sparse. (c) During the later stages of repopulation, the number of microglia and the spatial distribution return to baseline. (d-f) We then created and ran a mathematical model that sampled the number of microglia, (d) the persistent doublets, (e) the rapid divisions of microglia and (f) the secondary divisions of microglia during the peak of repopulation day 2-day 3. The mathematical model suggested that residual microglia can account for the rapid repopulation we observed *in vivo.*

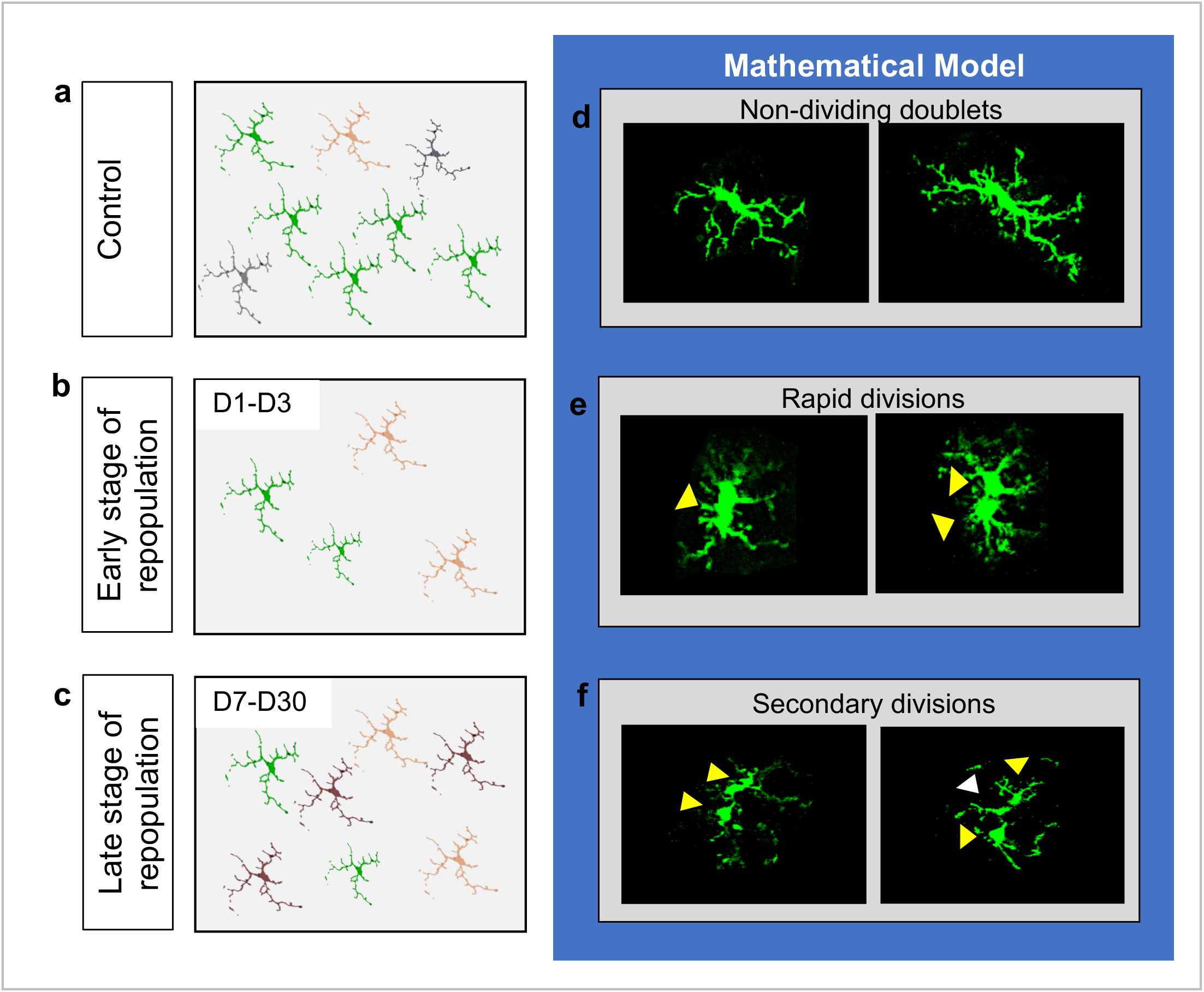

## Introduction

Microglia are the resident tissue macrophages of the brain (Crotti and Ransohoff, 2016; Ginhoux et al., 2010) with a developmental origin that is distinct from other macrophages (Ginhoux et al., 2010). In mice, microglial progenitor cells derive from the yolk sac and populate the brain before the blood brain barrier forms (Ginhoux et al., 2010; Kierdorf et al., 2013), slowly acquiring mature gene expression over a period of two to three weeks (Bennett et al., 2016). In both the rodent and human brain, microglial numbers are maintained throughout adult life (Fuger et al., 2017; Hashimoto et al., 2013; Reu et al., 2017), with no further contribution of peripheral cells to the microglial population in the absence of pathological changes (Ajami et al., 2007; Askew et al., 2017; Bruttger et al., 2015; Elmore et al., 2015; Mildner et al., 2007). Microglia are uniformly distributed in a distinct cellular grid throughout the brain parenchyma (Eyo et al., 2018; Hefendehl et al., 2014; Nimmerjahn et al., 2005) and maintain their territories with slow translocation on a timescale of days (Eyo et al., 2018).

Microglia are highly dynamic within their territory with extensive processes that make physical contact with cells in their environment (Bernier et al., 2019; Davalos, 2005; Nimmerjahn et al., 2005). Under basal conditions, motile microglial processes remodel, prune, and define neuronal circuits and establish transient physical contacts with synapses that influence neuronal circuits during early development and adulthood (Miyamoto, 2016; Paolicelli, 2011; Parkhurst et al., 2013; Schafer, 2010; Sipe et al., 2016; Tremblay et al., 2010). In addition to their many homeostatic roles in physiological brain function, these cells are traditionally known as a first line of defense in neuropathology and participate in responses to injury, infection, and neurological disease. For instance, chronic microglial activation is associated with a number of neurodegenerative and neurodevelopmental diseases such as Alzheimer’s Disease, Schizophrenia and Autism Spectrum Disorder (Bilbo et al., 2018; Kettenmann, 2011; Spittau, 2017).

While microglia are known to be long-lived cells, they do self-renew slowly and stochastically under unperturbed physiological conditions in the brain (Fuger et al., 2017). To study microglial self-renewal, recent studies have pharmacologically and genetically depleted microglia and uncovered a remarkable capacity for rapid repopulation of the microglial niche without a contribution of infiltrating peripheral monocytes (Ajami et al., 2007; Elmore et al., 2015). Early studies in fixed sections from mice following depletion suggested that microglial repopulation is driven by the proliferation of a specific nestin+ progenitor which divides in large proliferative macroclusters from which newly-born microglia migrate throughout the brain (Bruttger et al., 2015; Elmore et al., 2014). More recent studies have shown that newly-born microglia transiently express nestin early in repopulation; however, none of the repopulated microglia derive from nestin progenitors. Instead this static analysis of microglia in fixed tissue suggested that repopulated microglia are derived from the microglia that survive depletion in the cortex (<5%) rather than a specific progenitor population (Huang et al., 2018a; Huang et al., 2018b; Zhan et al., 2019). Similar to repopulation in the cortex, microglial repopulation in the retina was shown to be driven by remaining microglia; however, proliferation occurred in the central retina and these new microglia then migrated to repopulate peripheral areas, suggesting a specific site of microglial generation in the eye (Huang et al., 2018b; Zhang et al., 2018). In the retina, repopulated microglia eventually adopted the dynamic profiles of endogenous microglia before depletion, with similar morphologies, process dynamics and responses to focal injury 60 days following initial repopulation (Zhang et al., 2018), and repopulated cortical microglia also have expression patterns similar to those of endogenous microglia 30 days after repopulation (Bruttger et al., 2015; Huang et al., 2018b; Zhan et al., 2019). Genomic analysis of newly-born microglia in the cortex, however, show that new microglia within days of repopulation recapitulate developmental expression profiles (Elmore et al., 2015; Zhan et al., 2019), suggesting that newly-born microglia may undergo a stepwise maturation after repopulation that resembles developmental maturation.

While recent interest in microglia has illuminated many aspects of how microglia self-renew in the brain, many questions remain unanswered, including the site of microglial generation, the dynamics and loci of microglial proliferation, and the dynamics of how microglia acquire their mature characteristics once generated. It is critical to fully characterize the process of microglial self-renewal to understand how these cells, that are not generated in the yolk sac, differ from those that enter the brain early in development. Perturbations in microglial development can induce permanent changes in the microglial immune response (Bilbo et al., 2018; Knuesel et al., 2014), suggesting that microglia born in the adult brain, if they also undergo a period of maturation, may be particularly sensitive to homeostatic disturbance, affecting their function more so than that of long-lived microglia that originated from the yolk sac. Because microglia are critical to both physiological and pathological brain function, it is important to understand how these cells rearrange, renew, and mature as they are replenished in the mature brain.

Microglia are highly dynamic cells and understanding their behavior requires an equally dynamic approach that allows for monitoring their characteristics chronically over time in their native milieu. In our study, we aimed to characterize microglial ontogeny and maturation in the adult visual cortex using time-lapse imaging *in vivo* in awake young adult mice after microglial depletion. We show that microglial self-renewal is slow under basal conditions, with microglia maintaining their territories and showing little movement, loss or proliferation. In agreement with previous studies, we show that, following depletion, newly-born microglia rapidly repopulate the brain and acquire equal cell-to-cell spacing reminiscent of baseline conditions. Self-renewal is driven locally by residual microglia which have a capacity for fast and continued self-division that allows for rapid repopulation of the cortex. To determine whether other mechanisms such as migration from sites outside of the imaging area contributed to the increase in microglia numbers, we mathematically modeled microglial self-renewal *in vivo* and showed that the observed division rates could account for the rapid repopulation. Finally, we also showed that newly-born microglia very quickly acquire ramified morphologies as well as dynamic surveillance capabilities in response to focal injury, suggesting that microglial functional maturation is remarkably rapid in the adult brain and does not recapitulate developmental profiles. These findings suggest that the microglia landscape following depletion is restored through a rapid division of remaining microglia, local migration to fill the microglia niche and a fast acquisition of mature characteristics soon after repopulation.

## Results

### *In vivo* imaging of microglia shows limited migration and turnover with preserved territories in the physiological brain

To determine how microglia, self-renew, we first determined how much microglia movement and turnover occurred with our imaging approach in the absence of perturbation. To track microglial turnover *in vivo* under basal conditions, we imaged microglia daily in the same awake young adult mice using the microglial-labeled CX_3_Cr1^GFP/+^ transgenic mouse line (Jung et al., 2000) and a chronic cranial window preparation. Because a growing body of literature suggests that microglia behave differently under anesthesia (Li et al., 2012; Stowell et al., 2019; Sun et al., 2019) we wanted to capture the dynamics of microglia in the absence of anesthesia to avoid potentially inducing long-term alteration in microglial movement and turnover during repopulation. Microglia in the same cortical area could easily be imaged and tracked from day to day (Figure 1a and Supplementary video 1) without overt changes in microglial morphology which could indicate immune activation over time. Microglia numbers were stable over 14 consecutive days, and individual microglia could be re-identified in the same location daily over this period, suggesting limited local migration and a maintenance of microglial territories during this time (Figure 1a). Quantitative three-dimensional nearest neighbor (NN) analysis revealed that microglia generally maintain their own territories, whereby each microglia soma lies at ∼30 μm from its nearest neighboring microglia. This NN distance remained similar on subsequent days and between animals (Figure 1b, n=3, 16-17 microglia per mouse, 30-40 μm stacks). This is consistent with the classic distribution of microglia under physiological conditions whereby each microglia had its own region, with little overlap between neighboring territories, and suggests limited dynamic changes in distribution at baseline.

**Figure 1:**
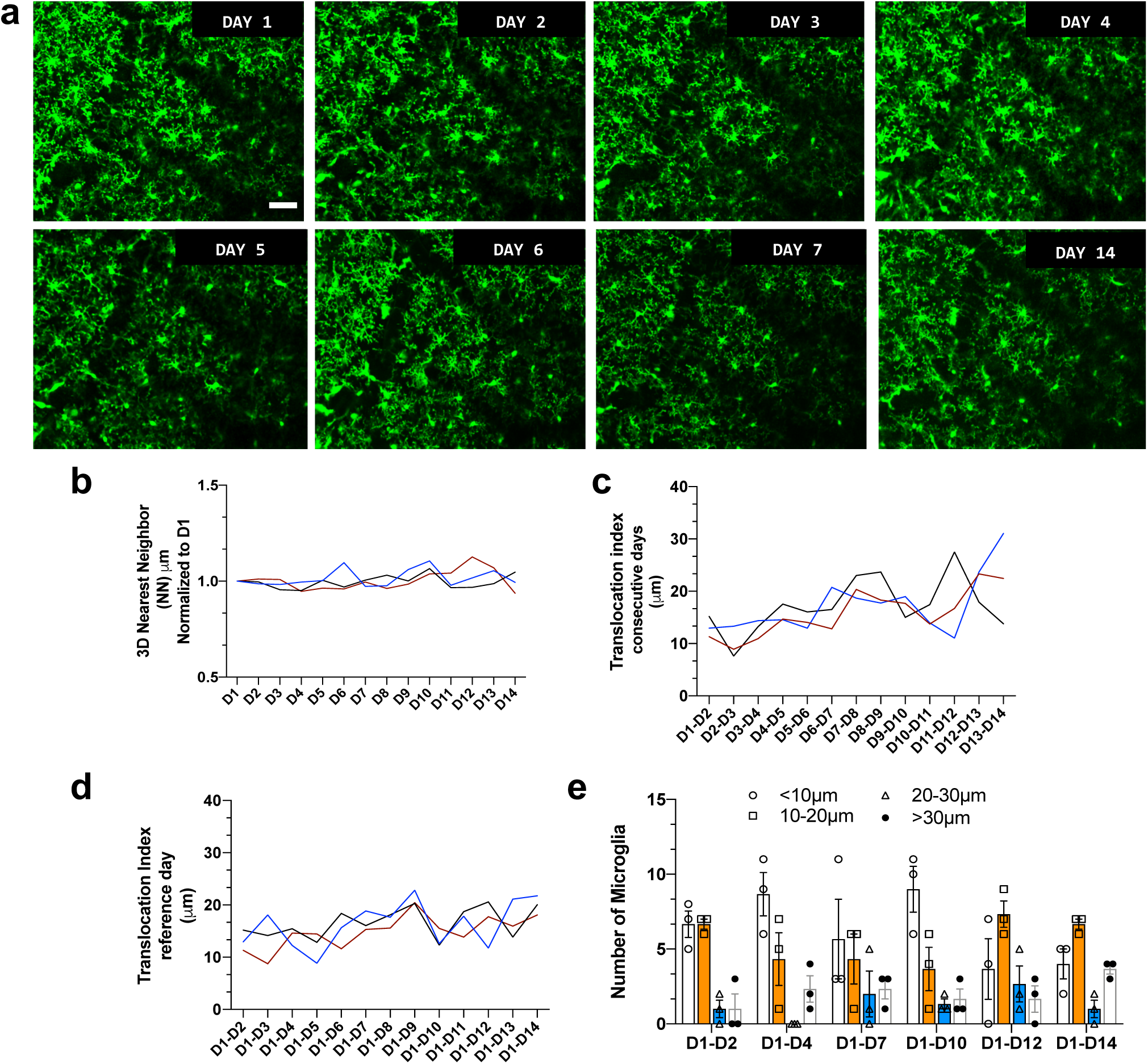
*In vivo* imaging of microglia shows limited migration and turnover in the physiological brain. (a) A field of microglia in an awake mouse imaged over 14 consecutive days. (b) Nearest neighbor quantification in 3D demonstrates the distribution of neighboring microglial cells over consecutive days. (c) The translocation index, which captured the average displacement of microglia over time, was ∼15um when consecutive imaging sessions were compared. (d) The translocation index increased when D1 is compared to imaging carried out later (D2-D14). (e) Microglia translocation between D1-D2, D1-D4, D1-D7, D1-D10, D1-D12 and D1-D14. On average, the majority of microglia remained within ∼10µm away of their original location. The proportion of microglia that moved within their domain (10-30 µm), (20-30 µm) or translocated a further distance (>30 µm) stayed relatively constant with increasing interval between imaging sessions (n=3, 30-40µm stacks, 13-17 microglia per mouse). Scale bar, 50*μ*m.

Microglial distribution over time was assessed with a custom algorithm which compared the location of microglia in the same field of view over the 14-day imaging period. To measure microglial movement, we defined a “translocation index”, which was an average distance between the location of each microglia on day 1 and the location of the nearest microglia on the n^th^ day (n=3, 16-17 microglia per mouse, 30-40 μm stacks). On average, microglia shifted on the order of 15 μm from day to day (Figure 1c), and this number increased as the two imaging sessions were further apart in time (Figure 1d), possibly due to tissue distortion over time. In general, most microglia did not move by more than 10 μm in either short (1-2 day), (1-4 day), (1-7 day) and long (1-10 day), (1-12 day), (1-14 day) comparisons. The proportion of microglia that moved more than 30 μm (the territory of a single microglia) was small in both short and long-term comparisons (Figure 1d-e, n=3, 16-17 microglia per mouse, 30-40 μm stacks). Overall, this analysis suggests that microglia are stable under basal conditions and show limited movement within their territory from over 14 days.

### Microglia rapidly replenish after partial depletion in the visual cortex

Microglial homeostasis is tightly regulated under basal conditions. Therefore, to explore microglia self-renewal in the visual cortex, we used an established paradigm which partially depletes microglia using PLX5622 (PLX), a Colony Stimulating Factor 1 Receptor (CSF1R/c-kit/Flt3) inhibitor (Dagher et al., 2015; Elmore et al., 2014; Najafi et al., 2018). With introduction of PLX for 7 days, microglia numbers in the visual cortex decreased by 70% (Figure 2a, b; Supplementary Figure 1) and the 3D NN number increased to 200 μm reflecting the decreased density and therefore increased distance between remaining microglia (Figure 2c). We did not observe changes in cytokine production or astrocyte morphology following depletion (Supplementary figures 2 and 3). The remaining 20-30% of microglia after depletion (7D PLX) and during the early stages of repopulation (D1) appeared to exhibit altered morphologies with two main groups represented: one with extensive processes and enlarged somas (yellow arrows) and the 2^nd^ with thickened and retracted processes (white arrows) (Figure 2a). Qualitatively, it appeared both morphological groups of remaining microglia were randomly organized suggesting that these morphologies were not driven by local cues.

**Figure 2:**
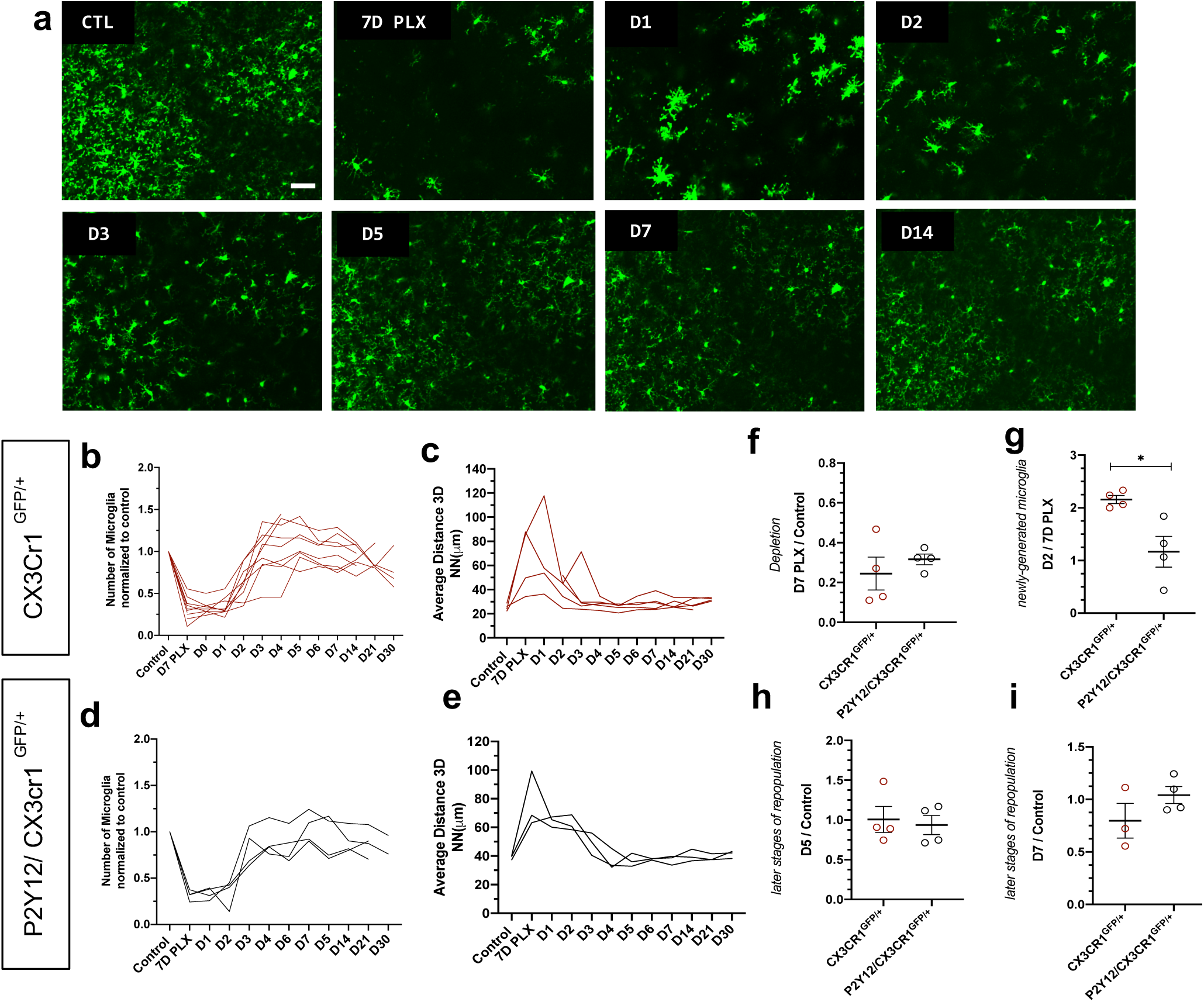
Microglia rapidly repopulate the visual cortex after partial depletion. (a) A field of microglia during depletion and repopulation imaged *in vivo* in the same awake mouse. (b) The number of microglia (normalized to control) during depletion (PLX) and with repopulation (day 1-day 30). Each line represents an individual animal (n=11, 100-150 microglia per mouse). (c) 3D nearest neighbor quantification showed a large increase during depletion and the early stages of repopulation before returning to control numbers (n=11, 100-150 microglia per mouse). (d) Depletion and repopulation dynamics were similar in the absence of P2Y12 (n=3-4 animals, 20-150 microglia per mouse). (e) 3D nearest neighbor analysis shows similar changes in microglial distribution during repopulation in the absence of P2Y12 (n=3 animals 20-150 microglia per mouse). (f) The ratio of microglia numbers observed on D7 PLX to control. (g) Repopulation was slightly delayed in P2Y12KO mice as compared WT, as the change in microglial numbers from depletion (PLX) to day 2 of repopulation was significantly smaller in the absence of P2Y12. (h) By day 5 of repopulation, the change in microglial numbers had normalized between WT and P2Y12KO mice. (i) Microglial numbers never fully recovered to control conditions in either WT or P2Y12KO mice. (f-i, t-test * p<0.05, n=4, 20-150 microglia per mouse). Scale bar, 50*μ*m.

After discontinuing PLX treatment, we imaged microglial restoration daily in the same mouse to capture the dynamics of this process (Figure 2a). Newly-born microglia rapidly repopulated the visual cortex and microglia numbers were almost completely restored after only 3 days of repopulation. In addition, the newly generated microglia surpassed baseline numbers after 5-7 days of repopulation (Figure 2b). In concert, the 3D NN distance returned to control levels after 3 days of repopulation and remained relatively stable until 30 days of repopulation, the last time point examined, at which time the number of microglia were similar to that of control microglia numbers before PLX treatment (Figure 2c and Supplementary video 2). These results suggest that microglial proliferation occurs very rapidly over a 24-hour period starting at ∼2 days after cessation of PLX treatment, and that microglia rapidly regain their territories within the visual cortex, maintaining their equal spacing and numbers after repopulation is complete.

To dissect the underlying mechanisms responsible for our observations of microglial repopulation, we considered signaling pathways in microglia that are associated with migration, microglial motility and maturation. While there are a number of receptors expressed by microglia that respond to changes in their environment, we focused on the P2Y12 receptor. P2Y12 is highly expressed exclusively in microglia in the brain (Bennett, 2016; Zhang et al., 2014) and regulates microglial translocation under physiological conditions *in vivo* (Eyo et al., 2014; Haynes et al., 2006). PLX treatment in P2Y12KO/CX_3_Cr1^GFP/+^ mice also caused a depletion of microglia of ∼70% (Figure 2d), with a concomitant increase in 3D NN (Figure 2e). The change in NN distance was not as profound as in CX_3_Cr1^GFP/+^ mice, possibly because these animals had, on average, smaller magnitudes of depletion, although depletion was highly variable across all animals examined irrespective of genotype (Figure 2f). While microglia repopulated in a temporal pattern similar to CX_3_Cr1^GFP/+^ mice after cessation of the inhibitor (Figure 2d, e), there appeared to be a slight delay in early repopulation (Figure 2g). However, this delay resolved quickly, and on day 5 and beyond microglial repopulation matched CX_3_Cr1^GFP/+^ controls (Figure 2h, i). This suggests that P2Y12 signaling plays a minor role in the repopulation dynamics of adult microglia.

### Residual microglia are capable of rapid division to generate new microglia and repopulate the cortex

Microglial self-renewal may be driven by local residual microglia that remain following depletion, although the dynamics of this process are poorly understood. It is unclear whether specific subpopulations of remaining microglia are responsible for division or all remaining cells have the capacity to divide; whether repopulation is local or originates in a specific brain region and is coupled to large scale microglial migration; and whether repopulating cells do so by single cell division or through an intermediate multinucleate body gives rise to multiple new microglia. During the repopulation phase, we observed occasional splitting of an existing microglia into two daughter cells, which has been reported previously (Fuger et al., 2017; Tay et al., 2017). In these cases, the appearance of new cells was associated with a characteristic increase and elongation of the cell soma of the original cell before it divided into two microglia (Figure 3a; Supplementary video 3). The original microglia often had extensive processes which were maintained during the division and generation of a new microglia (Figure 3a). Thus, newly generated microglia frequently had a ramified microglia arbor on the same day that division was complete. In many cases, we observed that newly born microglia had a long terminal process extending from one of the new cells (Figure 3a; top left panel inset). While one microglia cell remained stable within the network of cells and persisted at the imaged position over time, the other microglia moved away and took up new territory adjacent to the parent cell (Figure 3a bottom right panel; Supplementary video 3). Both cells were present in subsequent imaging sessions, indicating that these cells persist and become integrated in the microglial network.

**Figure 3:**
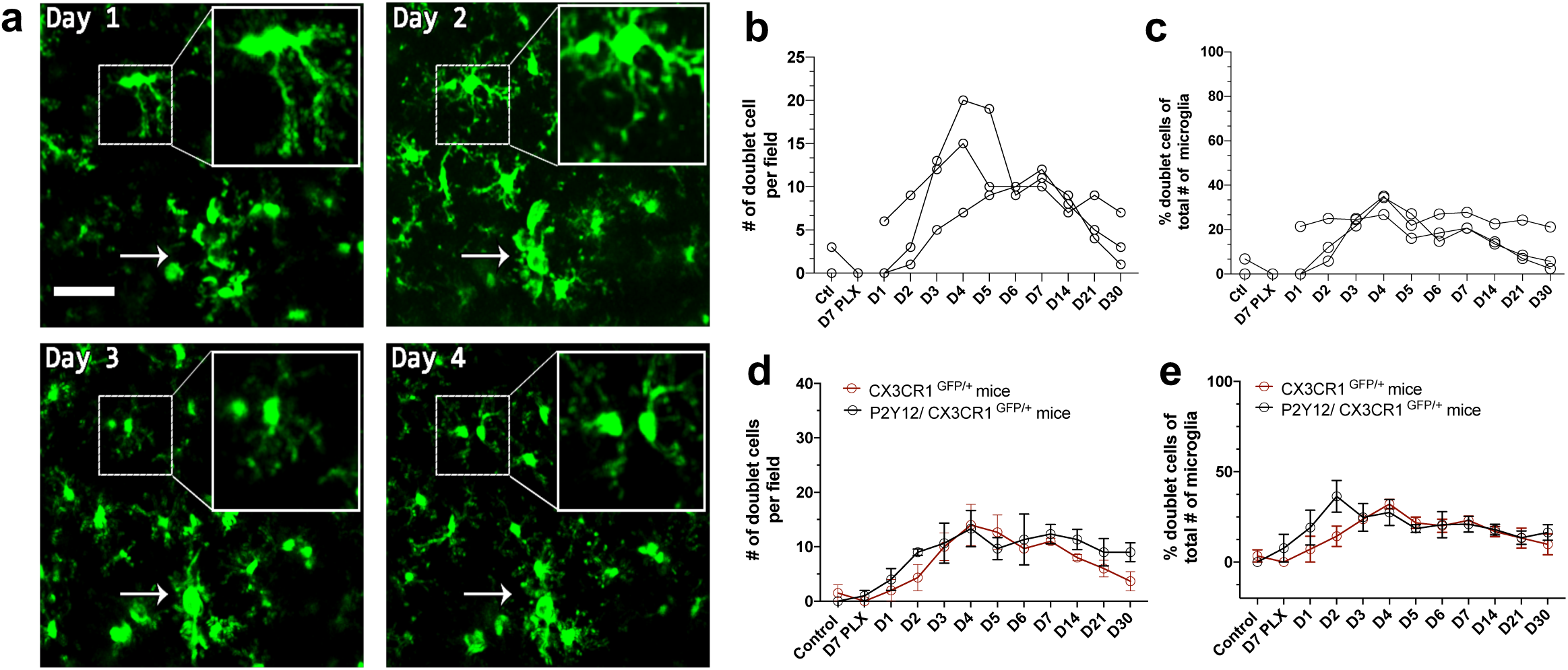
Microglia self-renewal via residual cells in the visual cortex. (a) Representative *in vivo* two photon imaging of microglial division in CX3CR1^GFP/+^ awake mice. Inset is a magnified image of the dividing doublet microglia, which appears as an elongated cell body at day 1 and divides into two cells at day 2. The two microglia then migrated away from one another over the subsequent 2 days. The arrow indicates a cluster of microglia that may represent a multinucleate body but did not generate new microglia during the imaging period. (b) The number of doublet cells per field increased at day 3 - day 5 of repopulation (c) Doublets made up close to 40% of the total number of microglia at day 4 of repopulation. (n=3-4, 50-150 microglia per mouse, 100µm stacks). The number (d) and percentage (e) of microglial doublets over time were similar in P2Y12KO mice as compared to WT (n=3 animals per group). Scale bar, 50µm. Lines represent individual animals.

The characteristic morphologies of microglia in the process of division allowed us to identify “potential” proliferating cells, which we refer to as doublet somas. We quantified both the total number of doublets and the percentage of all microglia that were doublets during our imaging of depletion and repopulation and found that both these measures were increased in the repopulation phase (Figure 3b, c). While these dividing cells could contribute to the rapid increase in microglial numbers observed, two pieces of evidence suggested that other mechanisms may be important. First, in our model of depletion, the number of doublet cells per field increased around day 3 to day 5 of repopulation but microglia had largely repopulated the brain by day 3 after cessation of PLX. Second, the time scale of division was slow (Figure 2b and 3b, c). The percentage of microglia that were in a doublet state was also low even during peak repopulation, and this level of doublets in the microglial population was maintained long after microglial repopulation was complete (Figure 2b and 3b, c). A similar temporal profile of doublet microglia was observed in P2Y12KO/CX_3_Cr1^GFP/+^ mice, again suggesting that P2Y12 signaling did not play a large role in regulating microglial division (Figure 3d, e). Secondly, the time scale of division was slow and the same microglia doublet could be observed for many days before a split was first evident (Figure 3a), with spatial separation of the daughter cells similarly taking multiple days. Thus, the slow rate of division is unlikely to allow these cells to completely repopulate the entire brain parenchyma rapidly after the cessation of PLX treatment.

### Mathematical modeling of microglial repopulation kinetics in the adult visual cortex

To determine whether microglia are capable of more rapid division that could account for the rapid increase in microglial numbers seen 2-3 days following cessation of PLX treatment, we imaged awake young adult mice every 4 hours for 24 hours during the most dynamic time of repopulation (Day 2-Day 3; Figure 4). Unlike what we observed during once per day imaging sessions (Figure 2 and Figure 3), microglial division could be remarkably rapid where doublets separated into two daughter cells in as little as 4 hours (Figure 4b). The rates of microglia division were not uniform and some cells divided slower within 8 or 12 hours or remained in the doublet state for the duration of imaging (Figure 4a). Notably, in some cases, we observed cells dividing twice (Figure 4c, d). The first division occurred rapidly, typically within 4 hours and the cells migrated away from one another. These cells then adopted a doublet morphology and divided again in less than 8 hours (Figure 4c, d). The rate of microglial division did not appear to be spatially regulated as microglia in close proximity divided at different times (Figure 4d). Microglial numbers roughly doubled over this imaging period in the majority of our animals (Figure 4e), and 3D nearest neighbor numbers fell as microglia were generated (Figure 4f). The number of doublet cells also increased but tended to plateau around 12 hours into the imaging period (Figure 4g), as did the number of total and primary divisions (Figure 4h). On the whole, dividing cells represented a subset of cells with a doublet morphology (Figure 4i) but the majority of cells with this morphology divided by the end of the imaging period. In fact, 50% of microglia present in the image during the first imaging session divided after 24 hours and the majority of non-dividing cells had a non-doublet morphology. Overall, this points to a remarkable capacity of the majority of the remaining microglia to divide rapidly and suggests that this rapid division may explain the fast repopulation of the cortex after depletion.

**Figure 4:**
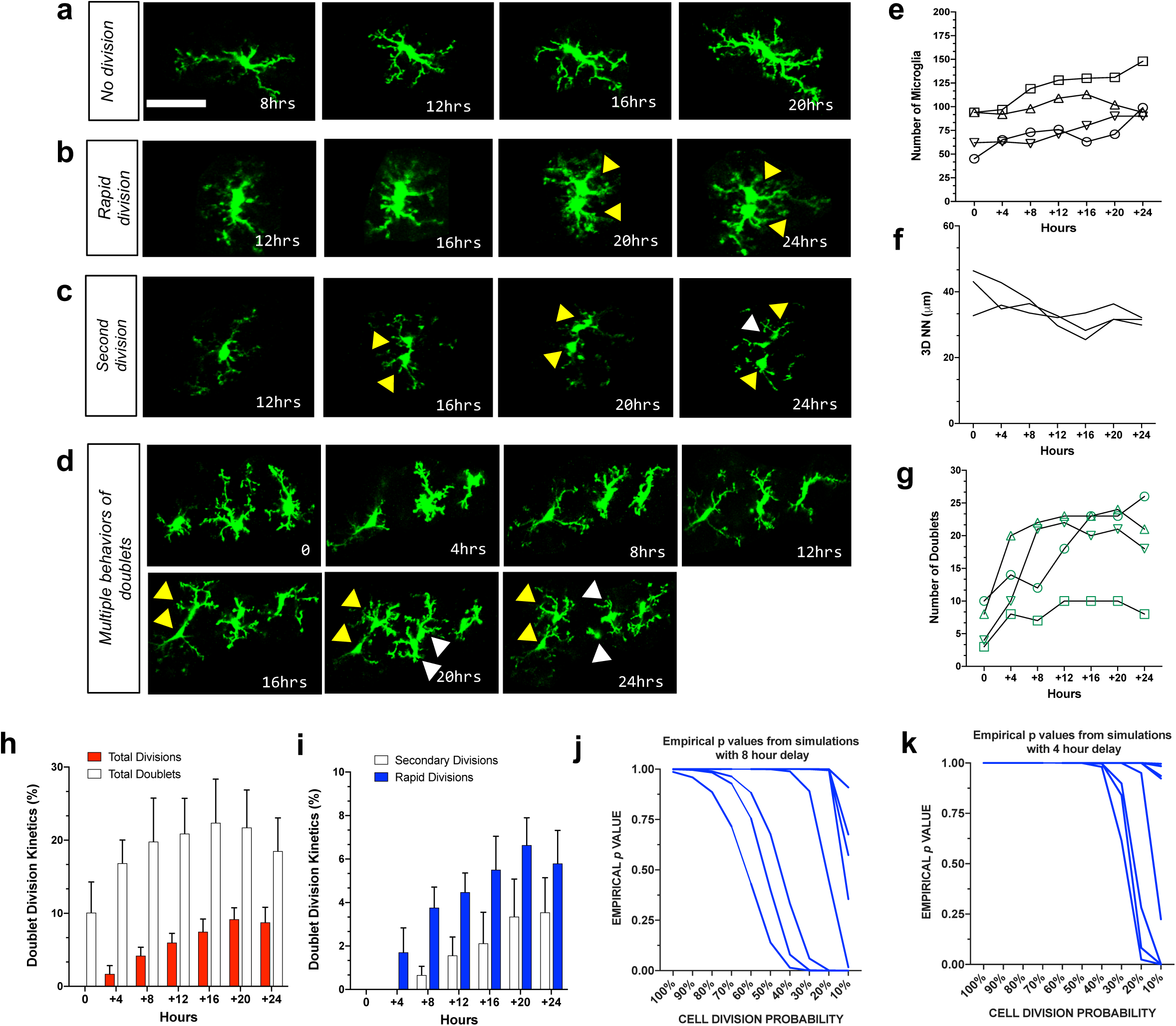
Mathematical modeling of microglia repopulation in the visual cortex demonstrate that cell division can account for the fast repopulation observed *in vivo.* We imaged CX3CR1^GFP/+^ mice every 4 hours for 24 hours during the peak of repopulation (day 2-day 3). We observed a range of behaviors of existing microglia: (a) no division (b) rapid division on the time scale of 4-8 hours once the doublet appeared and (c) secondary division, where newly divided microglia underwent another division. (d) Microglial divisions are not spatially regulated. Three microglia in the field of view divided at different times over the course of 24h. (e) The number of microglia during the 24h period increased in 3 of the 4 animals imaged. (f) Microglial 3D nearest neighbor distance decreased with time as more microglia were added to the population (n=3). (g) The number of microglial doublets increased over the 24h period. (h) Rapid divisions made up the larger proportion of divisions at each imaging time point compared to secondary divisions. (i) The proportion of microglia in the doublet state grew during this time, but on average doublets made up 20% of the total microglia. (j-k) Mathematical modeling of microglial repopulation using measured kinetic parameters. Empirical ρ values from the simulation are plotted for a minimum 4-hour delay (j) and 8-hour delay (k) between subsequent divisions. (j-k) 11 animals that were imaged daily (Fig. 2-3) were modeled and the simulation for each of these animals is represented by a line. (e-i) n=4 animals were imaged for 24hours during the peak of repopulation. Scale bar, 50*μ*m.

To determine whether local remaining microglia could repopulate the cortex through rapid division, we created a mathematical model which used the division dynamics quantified in the 24-hour imaging experiment to determine whether these rapid divisions could account for the increases in microglial numbers seen when repopulation was imaged daily (Figure 4j-k and Supplementary Figure 4). The model considered three parameters: (1) the number of cells in the population on day 2 of the daily imaging paradigm, (2) the probability that a cell will be eligible for division (we tested using proportions ranging from 10-100% in increments of 10), and (3) the empirical division rates of the cells during the peak of repopulation (Figure 4i). The simulation was run 500,000 times. We randomly sampled the number of cells in the population that are eligible for division from a binomial distribution with probability of success equal to the proportion of cells eligible for division and set that equal to the initial day. We counted the number of cells after 24 hours that divided, as well as all the cells in the process of dividing to account for the total number of cells in the final subtotal. In 4-hour intervals, we resampled cells from the population of those eligible for division (total number of cells in circulation minus cells previously selected for division) and corresponding division rates before concluding after 24 hours have passed. The empirical ρ -values from the simulation with a 4 and 8-hour delay between divisions suggest that it is reasonable to believe that the repopulation we observed from days 2 to 3 in our daily imaging experiments in 11 animals (Figure 2) came purely from local doublet cell division, as long as 50% of cells are capable of division (which is the number of dividing cells we observed when imaging every 4 hours; Figure 4j-k). In fact, in half of the animals imaged, only 10% of cells would need to be capable of division to repopulate the visual cortex. This data suggests that local division of remaining microglia is likely responsible for rapid repopulation with newly born microglia capable of further division and without the need for the migration of newly generated microglia from a different site of generation.

### Newly-born microglia acquire a hyperramified morphology in the later stages of repopulation

*In vivo* tracking and mathematical modeling of microglial self-renewal in the adult visual cortex show that repopulation is largely driven by remaining cells that form doublets and rapidly divide to repopulate the brain. However, it is still unclear how newly-born microglia mature in the visual cortex. We observed that doublet cells were ramified during division, resulting in new cells that also had ramified morphologies soon after they separated. Because microglial morphology is difficult to quantify in the wide-field imaging in awake animals, we imaged a subset of microglia during repopulation under anesthesia at high digital zoom in order to closely quantify the subtle changes in repopulated microglial morphology (Figure 5a). We then traced individual microglia and used Sholl analysis to assay the complexity of the arbors (Figure 5b and Supplementary Figure 5). At the peak of depletion (7 days of PLX), the remaining microglia exhibited a ramified morphology with extensive processes similar to the basal microglia process ramification under steady-state conditions (Figure 5b-d). In addition, these microglia have enlarged somas (Figure 5e). In the early phase of repopulation (day 1 – day 3), the newly-born microglia appear ramified with secondary processes (Figure 5b-d). After this, a more complex arbor begins to form (Figure 5b-d), whereby there is an overshoot in the maximum number of intersections and integrated area under the Sholl curve at 5 days of repopulation, which correlates with our increase in the number of microglia in our original repopulation data (Figure 3b and 5c-d). This slight hyperramification is maintained for as long as we imaged (30 days of repopulation; Figure 5c-d), despite a return to baseline numbers of microglia (Figure 3b) and a return to a smaller soma size (Figure 5e). Therefore, newly-born microglia rapidly acquire complex morphologies but exhibit a chronic hyperramification that distinguishes them from microglia in control conditions.

**Figure 5:**
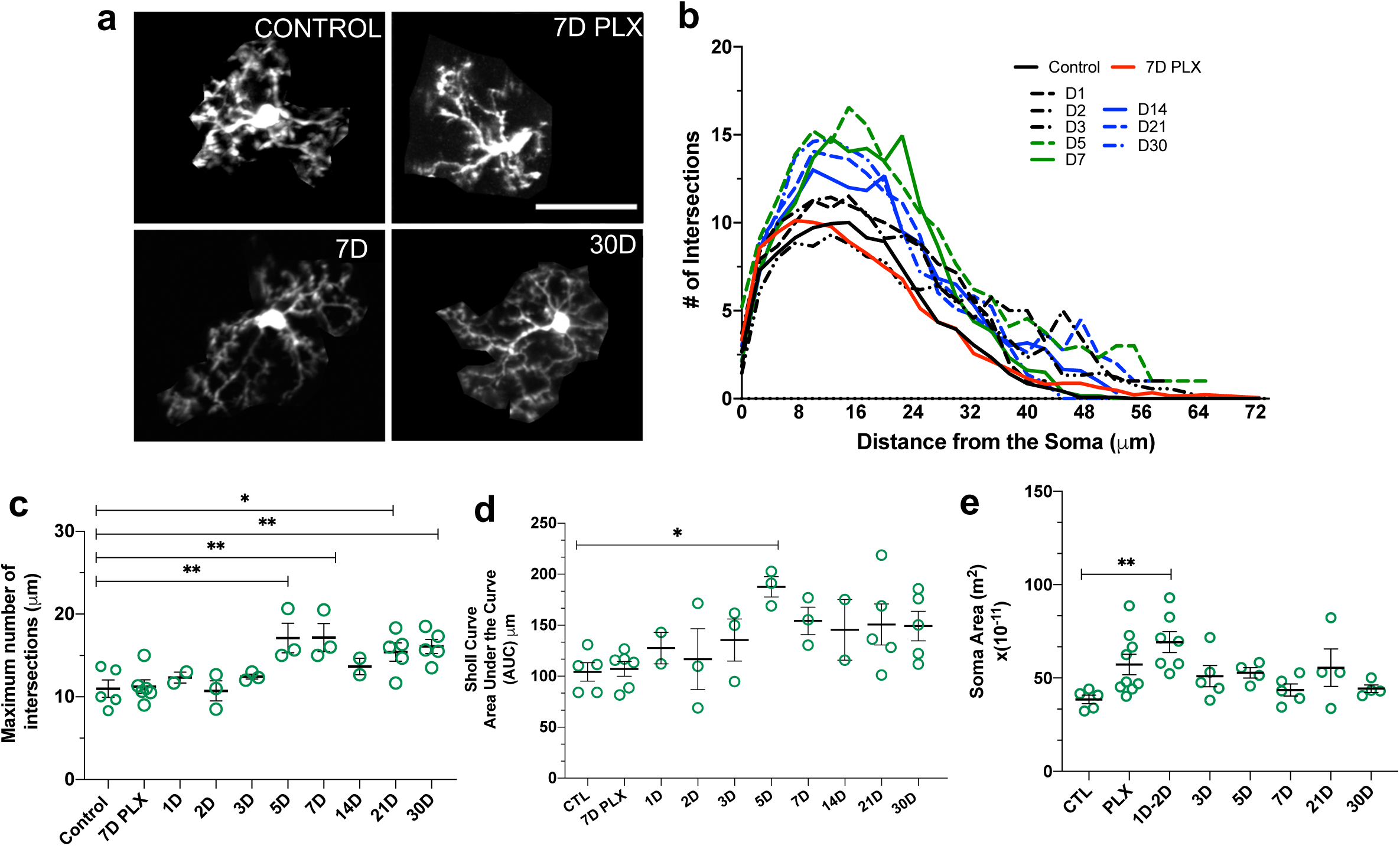
Newly-born microglia remain in a hyperramified state after repopulation. (a) Individual microglia imaged in mice during control, depletion (7 days of PLX) and repopulation (7 and 30 days). (b) Sholl profiles summarizing the morphology of microglia during depletion and repopulation. (c) Microglia have an increased number of maximum intersections in the later time points of repopulation; note the increased arborization after day 5 (n=3-5 animals per group, 2-4 microglia per mouse, *p<0.05, **p<0.01, One-way ANOVA, Dunnett post-hoc test). (d) Microglia have a greater AUC (Area under the curve) in the later stages of repopulation (n=3-5 animals per group, 2-4 microglia per mouse, *p<0.05, One-way ANOVA, Dunnett post-hoc test). (e) Microglia demonstrated a greater soma area during the early stages for repopulation (n=5-9 animals per group, n=3-6 microglia per mouse, **p<0.01, One-way ANOVA, Dunnett post-hoc test). Graphs show mean ± s.e.m. Points represent individual animals. Scale bar, 50*μ*m.

### Newly-born microglia are dynamic and survey the brain

While a handful of previous studies characterized newly-born microglia repopulation using a static approach, we chose to image microglia *in vivo* because these cells are highly motile allowing them to rapidly survey the parenchyma and perform homeostatic functions in the brain. Given the changes in microglial morphology after repopulation (Figure 5), we set out to determine if the differences were also reflected in their dynamics (Figure 6). Motility measurements were carried out under anesthesia as adrenergic signaling in the awake condition dampens microglial dynamics (Liu et al., 2019; Stowell et al., 2019; Sun et al., 2019), allowing us to reliably track microglial process extension and retraction on the order of minutes (Figure 6a-c). PLX treatment led to only a modest decrease in motility (Figure 6d), as imaged over a one-hour period, suggesting that the 20-30% of microglia that remain after depletion, not only have full arbors, but are motile and can effectively interact with elements in the parenchyma. This decrease in motility was accompanied by a decrease in instability, which measures the retraction of stable processes, (Figure 6e) but no change in the stability index, which measures the stabilization of newly extended processes (Figure 6f). As new microglia were generated after depletion, microglial motility recovered such that the increase in movement correlated with the increase in the number of microglia over time. In addition, during the later stages of repopulation (day 21-day 30), we found no significant difference in basal microglia process motility and long-term stability as compared to baseline (Figure 6d-f). Combined with our previous data on microglia repopulation, these data suggest that repopulation can be divided in two stages: early and late repopulation. As microglia mature in the later stages of repopulation (day 7 – day 30), they exhibit more complex arbors suggesting a dysregulation in their morphology. This long-term change in morphology, however, does not impact microglial motility which resembles basal microglial states (Figure 6d). Finally, to test whether changes in morphology and dynamics of new microglia affect microglial surveillance, we assessed microglial process coverage over one hour (Figure 6g-i). Following 7 days of PLX, the remaining microglia population survey a much smaller area than observed under basal conditions due to their reduced numbers. Microglial surveillance recovers at day 3 of repopulation when microglial numbers return to baseline values and is maintained until day 30 (Figure 6h). When surveillance was normalized to the original microglial morphology to account for the loss of microglia with depletion, we found that individual microglia surveilled their territory in the same manner throughout control, depletion and repopulation conditions (Figure 6i). Based on these findings, we conclude that newly-born microglia mature within days of repopulation, acquire complex arbors, and survey the environment effectively.

**Figure 6:**
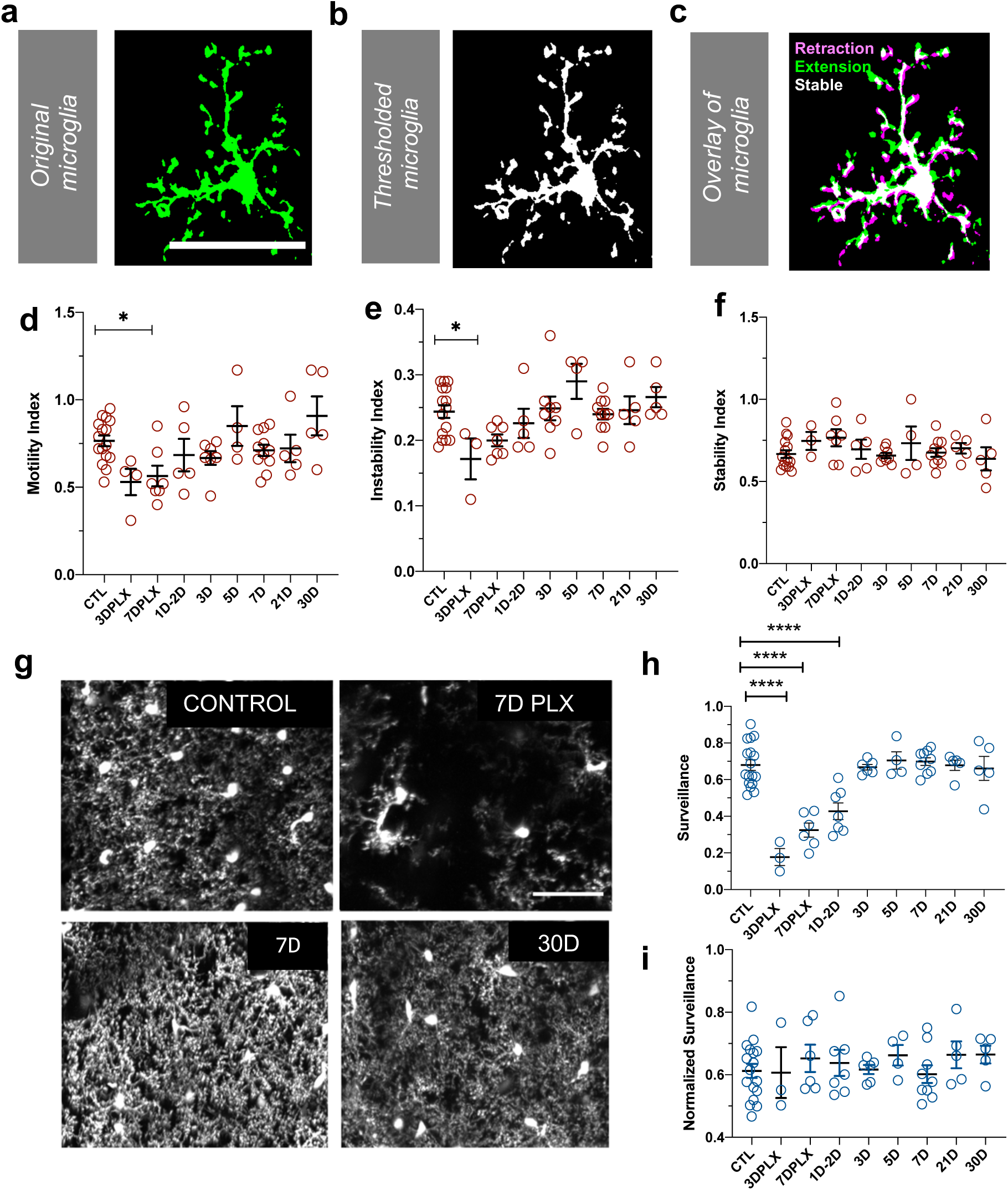
Newly-born microglia are dynamic and survey the brain. (a) Example of motility analysis showing the original microglia image, (b) Thresholded microglia (c) and a visual representation of microglial motility whereby thresholded imaged from two time points are combined with magenta representing retraction, green representing extension and white representing stable pixels. (d) Quantification of the motility index which compared the gain and loss of pixels across 5-minute intervals. Microglia motility was decreased during depletion but recovered quickly with repopulation. (e) A similar trend was observed in the instability index which was calculated as the proportion of stable pixels that became retracted over the total stable pixels in the first time point. (f) No change in the stability index (the proportion of extended pixels that became stable divided by the total extended pixels in the first overlay) was observed. (g) Maximum projection of microglial processes over the hour imaging session to observe surveillance (control, 7 days of PLX, 7 days of repopulation and 30 days of repopulation). (h) Microglial surveillance of the parenchyma dropped as microglia were depleted from the brain, but recovered quickly during repopulation as soon as microglial numbers reached control levels (3 day). (i) Graph of surveillance normalized to the extent of microglial coverage in the first time point. (n=3-14 animals per group, *p<0.05, ****p<0.0001, One-way ANOVA, Dunnett post-hoc test). Graphs show mean ± s.e.m. Points represent individual animals. Scale bar, 50µm.

### Newly-born microglia respond robustly to acute focal tissue injury

To determine whether newly-born microglia can carry out their normal pathological functions, we generated focal laser ablation injuries in the visual cortex using the two-photon laser microscope and quantified the movement of microglial processes toward the site of injury over the course of 1 hour using two separate methods (Figure 7 and Supplementary video 4). First, we developed a custom optic flow-based algorithm to calculate the directional velocity of microglia processes moving toward the injury site (Figure 7e-f). Analysis of the average of all vectors moving towards the core showed that microglia at all stages of depletion and repopulation responded robustly to laser ablation injury (Figure 7f, g). Both the maximum magnitude and the integrated response (area under the curve) were similar across conditions (Figure 7g, h). It is interesting to note that, under PLX treatment, the remaining microglia, although less motile, sparse and randomly organized in the cortex, responded robustly to laser ablation injury with only a slight trend towards a decreased response (Figure 7f, g, h). A similar pattern was observed with convergence analysis which calculates the number of pixels representing microglia processes entering a small ring around the core of the injury site (Figure 7i). Similar to the directional velocity measure, there was no difference in the maximum convergence at 60 minutes and the area under the curve value at each time point (Figure 7j, k). This suggests that the newly-born microglia during the early phases of repopulation are capable of responding robustly to focal laser ablation injury. This is an indication that microglia retain the necessary sensors such as P2Y12 and internal machinery to respond to injury.

**Figure 7:**
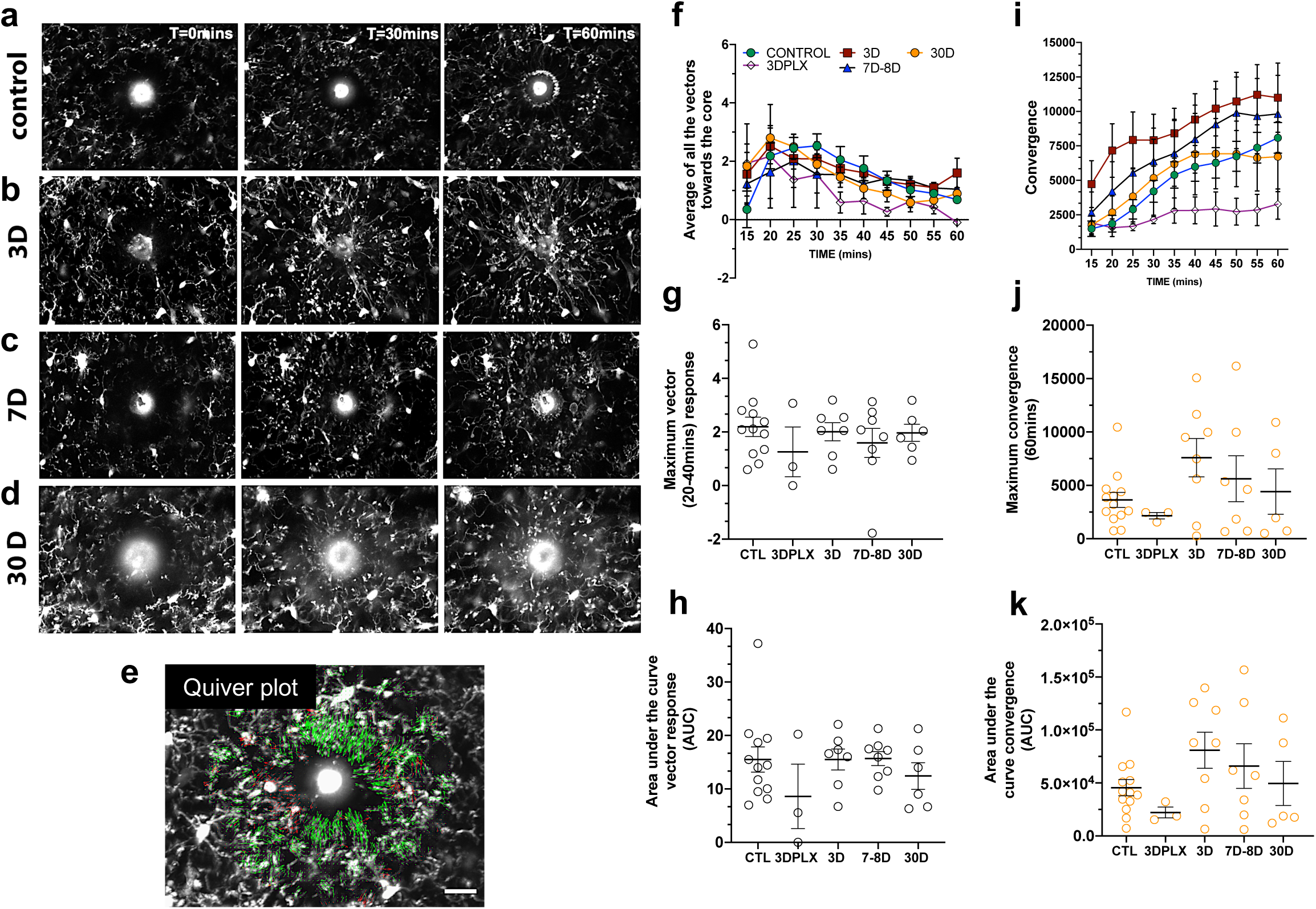
Newly-born microglia respond robustly to acute laser ablation injury. Example microglial response to focal laser ablation at t=0 min, t=30min and t=55 min after ablation for control (a), 3 days of repopulation (b), 7 days of repopulation (c), and 30 days of repopulation (d). (e) Quiver plot of microglial response. Green arrows correspond to vectors moving towards the core and red arrows correspond to vectors moving away from the core. (f) Graph showing the average vectors moving towards the core following focal laser ablation injury. (g-h) There were no statistically significant differences observed in the maximum directional response, or the integrated area under the curve over time. The dynamics of the convergence of microglial processes on the injury core (n=12 (control), n=3 (3 days PLX), n=8 (7-8 days), n=6 (30 days). (i) Graph showing the convergence towards to core following focal laser ablation injury. (j-k There was no significant difference in the maximum convergence at 60 minutes, or the convergence response when assayed using the area under curve of the convergence graphs over time. n=3-12 animals per group, One-way ANOVA, Dunnett post hoc test. Graphs show mean ± s.e.m; ns. Points represent individual animals. Scale bar, 20µm.

## Discussion

In this study, we characterized in detail the dynamics of adult microglia ontogeny and maturation in the visual cortex of mice. Because microglia behave differently in awake vs. anesthetized states (Liu et al., 2019; Stowell et al., 2019; Sun et al., 2019), we used an awake *in vivo* chronic imaging setup to track microglial repopulation. First, we showed that microglia maintain their numbers and territories under basal conditions. Next, we found that microglia can divide rapidly and continuously to repopulate the brain within a few days after partial depletion with the colony stimulating factor 1 receptor (CSF1R) inhibitor, PLX, in a manner that is largely independent of signaling through the P2Y12R. A mathematical model based on the imaged kinetics of division confirmed that the repopulation was driven by very rapid, local self-renewal by surviving microglia which transition to a doublet morphology, divide and migrate apart before undergoing the next cycle of division. Finally, we found that newly-born microglia mature rapidly both structurally and functionally.

### The dynamics of microglial division are heterogeneous

Microglia are thought to be long-lived cells, self-renewing slowly and stochastically under unperturbed physiological conditions in the brain with an average lifetime of four years in human cortex (Reu et al., 2017), and 15 months in the mouse (Fuger et al., 2017). Despite their stability, microglia can rapidly repopulate the brain when their niche has been depleted (Bruttger et al., 2015; Elmore et al., 2015). While this remarkable capacity for self-renewal has been the subject of intense interest, many questions remain unanswered, in particular regarding the kinetics of microglia self-renewal. By taking a chronic *in vivo* approach coupled with mathematical modeling, we were able to identify and track novel behaviors of residual and dividing microglia *in vivo*. Based on the average cell numbers and the various cell division rates, our simulations of microglia repopulation support the conclusion that the residual cells that remain after depletion can rapidly divide to repopulate the visual cortex.

We observed that during the height of repopulation, ∼50% of cells adopted a doublet morphology indicative of an impending division, suggesting that most residual cells are capable of proliferating. While we also observed these doublet morphologies after repopulation was complete, the division rates at that time were slow with division occurring over a period of days, as seen in the non-depleted brain (Fuger et al., 2017), and inconsistent with rapid repopulation. In contrast, at the height of repopulation, division could occur very rapidly, with a cell adopting a doublet morphology and undergoing division within 4 hours. These rapidly dividing cells can undergo a secondary division within our 24-imaging period. This secondary division could occur in both cells generated from the first division, suggesting a remarkable capacity for proliferation in residual microglia.

However, the rates of microglia division were not uniform. While some cells divided rapidly and repeatedly, others divided slowly within 8 or 12 hours or remained in the doublet state for the 24-hour period of imaging. While we cannot discern whether these differences in division rates are tied to specific subpopulations of microglia, it is well known that microglia are morphologically and transcriptionally diverse, with high throughput single-cell transcriptomics identifying distinct subpopulations of microglia with unique molecular signatures that change with age and in response to insults (Ayata et al., 2018; Hammond et al., 2019; Li et al., 2019). This heterogeneity in microglia is also seen across various brain regions (Ayata et al., 2018; Hammond et al., 2019; O’Koren et al., 2019). Similar heterogeneity in the residual microglial population may play a role in the dynamics of repopulation, contributing to a heterogeneity in the capacity to divide. For instance, it has been suggested that the remaining microglia after depletion are a specialized CSF1R-inhibitor resistant population of Mac2-progenitor-like microglia (Zhan et., al 2019, BioRXiV). Thus, rapidly dividing cells may correspond to this specialized population which could carry the bulk of repopulation with slow contribution from other populations that survive depletion. A similar scenario has been observed in the retina where microglia repopulate from a distinct source and migrate through the optic nerve to repopulate the rest of the retina (Huang et al., 2018a). While more detailed studies will be needed to determine the extent of microglial heterogeneity after PLX treatment, characterizing the heterogeneity of repopulating microglia may provide new avenues for understanding the phenotypes of adult-generated vs. yolk sac born microglia.

### Newly generated microglia rapidly recapitulate functional features of endogenous microglia

We show that newly-born microglia acquire mature characteristics such as baseline motility, surveillance and response to acute laser ablation remarkably quickly. In fact, dividing doublet microglia have an extensive, motile arbor which is maintained through the division such that the two resulting cells already have extensive processes which surveil the environment and can respond to injury. While microglial expression in the early phase of development appears to recapitulate developmental programs (Zhan et al., 2019), we did not observe the typical morphologies of developing microglia, or changes in process dynamics and responses to injury that could indicate a less mature phenotype. This suggests that if microglia do revert to a developmental program when they are first generated in the adult brain, this profile does not significantly alter their physiological functions. Alternately, they may go through an accelerated developmental program, as expression also matures much faster than the stepwise development of microglia over a period of weeks in development (Bennett, 2016; Matcovitch-Natan et al., 2016; Zhan et al., 2019).

While many studies have shown that after 30 days of repopulation, new microglia largely adopt the gene expression profile of control microglia (Elmore et al., 2015; Zhan et al., 2019) several observations in our study suggest that adult born microglia may not be identical to their yolk sac-generated counterparts. We found that after repopulation, microglial structure was more complex, and that more microglia adopted a doublet-like morphology with an elongated cell body. Changes in morphology of microglia are traditionally associated with inflammation, with hypo-ramified reactive microglia being commonly associated with disease where they secrete pro-inflammatory factors (Fontainhas et al., 2011; Smith et al., 2019; Streit et al., 1999; Tanaka and Maeda, 1996). The hyperramified phenotype of newly-born microglia can be associated with a transient inflammatory activation but has also been observed in non-pathological, experience-dependent changes (Sipe et al., 2016; Tremblay et al., 2010). This persistent hyperramification could therefore be a change elicited by different genetic programs in the newly born microglia that alter their immune functions. While we did not see any evidence inflammatory signaling after depletion with PLX (Supplementary Figure 2), it is also possible that newly born microglia change their morphology in response to changes in the extracellular environment either due to damage elicited by CSF1R blockade, a change in neuronal activity or persistent changes in the extracellular milieu.

The presence of a persistent increased number of microglia with elongated, doublet-like cell bodies, after repopulation, could be indicative of further morphological alteration of newly born microglia, but it could also suggest an increased capacity for division as similar doublets have been identified as a main source of dividing cells in other studies (Tay et al., 2017; Zhang et al., 2018). If Mac2+ microglia that remain after depletion (Zhan et., al 2019, BioRXiV) have increased capacity for division, they may divide to produce new microglia that are distinct from yolk sac-generated microglia and rapidly take on mature characteristics, contributing to a different phenotype of adult vs. yolk-sac born microglia. The Mac2+ cells may remain in the population as persistent doublet cells primed for division and self-renewal. Thus, while adult born microglia appear to rapidly take on their mature roles in the brain, their characteristics may differ in subtle ways from microglia that are generated in the yolk sac and populate the brain in development.

### Newly generated microglia chronological versus biological age

Functional characterization of newly-born microglia may provide valuable clues for determining treatments for neurodegenerative and neurodevelopmental diseases where microglia are often thought to be dysregulated (Keren-Shaul et al., 2017; Krasemann et al., 2017; Perry et al., 2010). It is possible that efficient rejuvenation of old “senescent” microglia in the diseased brain with new microglia through depletion and repopulation can generate a new population of microglia with improved functions (Elmore et al., 2018; Rice et al., 2015; Spangenberg et al., 2016). However, it is important to recognize that it remains unclear whether the chronologically younger newly-born cell is also biologically younger. In fact, we do not yet understand the biological process of microglia repopulation and it is not known whether the division of microglia is symmetrical or asymmetrical. To determine whether a newly divided cell is in fact “rejuvenated”, the “maturity” of that cell will need to be tested directly. This could be assessed by looking at transcriptional and epigenetic signatures and comparing to similar studies done in development (Hammond et al., 2019; Matcovitch-Natan et al., 2016). Newly-born microglia may go through a stepwise expression program that partly recapitulates development giving them a younger functional profile, or they may acquire all the epigenetic and cytoplasmic features of their “aged” parent cell, leading to repopulation but not “rejuvenation” of the microglia niche (Zhan et al., 2019). This latter option could also explain why the newly-born cells acquire mature characteristics and functions early on in the repopulation phase.

### Intrinsic and extrinsic mechanisms of microglia repopulation

A combination of exogenous and endogenous signaling mechanisms may contribute to microglia rapid repopulation and spatial organization. Recent evidence has shed light onto some of the extracellular signals that shape repopulation, including those that regulate microglia proliferation such as colony stimulating factor 1 (CSF-1) and interleukin 1 receptor (IL1R), and those that regulate migration such as fractalkine through its receptor, CX3CR1 (Bruttger et al., 2015; Elmore et al., 2015; Zhang et al., 2018). A gradient or threshold of these factors may start intracellular cascades in microglia that trigger division and migration, such as NF-KB signaling which has been shown to be important for microglia to repopulate fully (Zhan et al., 2019). In addition, we show that the P2Y12 receptor is not critical to doublet formation, repopulation, or the acquisition of microglial territories (as reflected by changes in nearest neighbor quantification) in the visual cortex (Figure 2 and 3), which is surprising given that P2Y12 modulates microglial translocation in baseline conditions (Eyo et al., 2018). The progressive distribution of microglia to achieve tiling during repopulation is most likely maintained by lateral inhibition mechanisms and homeostatic microglia-specific genes such as Sal1 and Mafb—which both maintain microglia in a ramified nonclustered state (Buttgereit et al., 2016; Matcovitch-Natan et al., 2016). Because microglial numbers overshoot their target during the early stages of repopulation, it is also likely that astrocytes may be recruited to phagocytose excess newly-born microglia, and these cells may contribute to the extracellular environment that promotes microglial migration and maturation. Altogether, an interplay of diverse factors likely contributes to microglia homeostasis following depletion, which may include endogenous mechanisms at play in specific populations of microglia, including Mac2+ positive cells, as well as exogenous factors that come from microglia, neurons and astrocytes and work together to re-establish microglia numbers in the adult brain.

### Concluding remarks

Together our results build on current adult microglia ontogeny research showing that rejuvenation of adult microglia is driven by stochastic doublet cell division locally during repopulation, with newly born cells rapidly able to take on their roles in the brain. In addition, our mathematical modeling supports the idea that division of residual microglia alone can fill the microglia niche following depletion. While our *in vivo* tracking approach illuminates the rapid and heterogeneous dynamics of microglial repopulation, future studies will be needed to fully understand the mechanisms that lead to the generation of new microglia and their ability to adopt their mature features. Such studies will provide a deeper understanding into the important role microglia play in homeostasis and disease.

## Methods

### Experimental Animals

All animal work was performed according to the approved guidelines from the University of Rochester, Institutional Animal Care and Use Committee and conformed to the National Institute of Health (NIH). Animals were housed in a 12-hour light/12-hour dark cycle with food ad libitum. 3-6-month-old male and female heterozygous (CX3CR1 ^GFP/+^) GFP reporter mice expressing GFP under control of the fractalkine receptor (CX3CR1) promoter (Jung et al., 2000) and P2Y12KO/CX3CR1 GFP mice were used for *in vivo* live imaging of visual cortex microglia repopulation and dynamics. All mice were derived from a C57/Bl6 background.

### Microglia Depletion

Diet (AIN-76A-D1001i, Research Diets, New Jersey, USA) containing 1200 mg/kg PLX5622 (Plexxikon Inc., Berkeley, CA, USA) was given to mice as the sole food source for 1-2 weeks to deplete microglia. Control diet with the same base formula but without the compound was given to the control group (Bl6 mice and YFP mice in Supplementary figure 1 and 2 that are not exposed to PLX) and during the repopulation phase for 3-4 weeks.

### Cranial Window Surgery

Animals were anesthetized using a fentanyl cocktail (i.p.) during the cranial window implantation surgical procedure. The fentanyl cocktail consisted of fentanyl (0.05 mg kg^-1^), midazolam (5.0 mg kg^-1^) and dexmedetomidine (0.5 mg kg^-1^). Body temperature was maintained at 37? C with a heating pad and aseptic technique was maintained during all surgical procedures. Mice were placed in a stereotaxic frame and head-fixed for cranial window surgeries. Hair was removed and the skull was exposed through a scalp incision. A 3mm Biopsy punch (Integra, Plainsboro, NJ, USA) was used to create a circular score on the skull over V1. A 0.5mm drill bit (FST, Foster City, CA) was used to then drill through the skull for the craniotomy, tracing the 3mm score. A 5mm coverslip attached to a 3mm coverslip (Warner Instruments, Harvard Bioscience, Hamden, CT) by UV glue (Norland Optical Adhesive, Norland Inc, Cranbury, NJ, USA) was then slowly lowered down into the craniotomy (3mm Side down). The coverslip was secured with Loctite 404 glue (Henkel Corp, Bridgewater, NJ, USA). A custom headplate produced by emachine shop (www.emachineshop.com) (designs courtesy of the Mriganka Sur Lab, MIT) was kept in place with C&B Metabond Dental cement (Parkell Inc, Brentwood, NY, USA). The dental cement was used to cover exposed skull and keep the headplate in place. Mice were administered slow-release buprenex by veterinary staff (s.q. mg kg-1 every 72 hours) and monitored for 72 hours.

### Two Photon Microscopy

A custom two-photon laser-scanning microscope was used for *in vivo* imaging (Ti: Sapphire, Mai-Tai, Spectraphysics; modified Fluoview confocal scan head, 20x water immersion objective lens, 0.95 numerical aperture, Olympus, Center Valley, PA). Excitation for fluorescent imaging was achieved with 100-fs laser pulses (80MHz) tuned to 920nm for GFP with a power of ∼30-40mW measured at the sample. Fluorescence was detected using a photomultiplier tube in whole-field detection mode using a 580/180 filter. Images were collected from 20μm -300μm into the brain. For repeated imaging, blood vessels were used as gross landmarks and stable microglia were also used as fine landmarks to reidentify the correct region for imaging. Image analysis was done offline using ImageJ and Matlab with custom algorithms as described in (Stowell et al., 2019) and available on request.

### Awake Imaging Sessions

During a two-week recovery from cranial window surgeries, mice were trained and habituated to awake imaging. Mice were head fixed under the microscope for 10 minutes on the first day and the time spent head fixed was increased by 5 minutes daily. Habituation sessions were terminated if mice exhibited signs of discomfort such as excessive movement and vocalization. Training was complete once mice tolerated 4 consecutive days of head fixation. Imaging session did not exceed 60 minutes daily.

### Microglia Migration Analysis

Two-photon XYZ images of microglia were collected. For analysis, microglia within a volume of 800 X 600 X 100 μm (approximately 60 to160 μm from the surface of the brain) were used. The images were processed and aligned using Fiji imageJ. The centroid for each microglia was identified and the x, y and z coordinates were recorded. Images were aligned over consecutive days using the blood vessels as gross landmarks. A custom algorithm was created to quantify the distances between individual microglia and the remaining microglia in the image. The minimum value was then used as the 3D nearest neighbor value for that microglia and nearest neighbor values were averaged together for all microglia in a single animal at each time point. Distances were calculated as shown below:

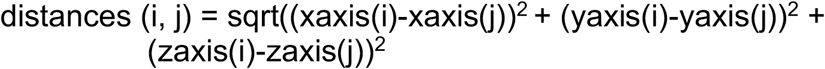

i, j corresponds to number of microglia in the matrix used to calculate the NN.

*Microglial Translocation:* The location of selected microglia from the first time point were identified and a the XYZ coordinates for the microglia were recorded for the first day. The 3D Nearest neighbor values were calculated as shown above except the location of a microglia was compared to the location of all microglia at another time point. Usually translocation was measured relative to Day 1 of imaging or the time point immediately prior.

### Mathematical modeling of microglial repopulation

We began with three parameters:

1. *N*_*0*_, the number of cells in the population on day 2 of the daily imaging paradigm,
2. *P*, a specified probability that a cell will undergo division,
3. *R*, the observed division rates in experimental data.

The algorithm was then initialized with the following values. The number of cells chosen to divide from *N*_*0*_ is given by _*0*_and were selected by sampling from a binomial distribution of size *N*_*0*_with probability *P*. Each cell selected for division was assigned a division rate by generating a sample *S*_*0*_ of size *D*_*0*_ from *R*. This process was repeated every 4 hours over a 24-hour period. We also considered cells to be ineligible for repeated division for a period of 4 hours after having divided. Using the specified notation and letting *i* = 1,2,3,4,5,6 and t_*n*_ = 4*n* we can denote the number of microglia in the population during the interval by *N*_*i*_, where

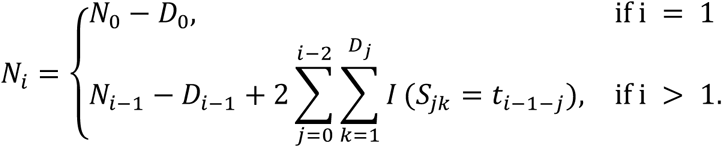

The final number of cells counted in the population were given by the sum total of cells that have completed division and those that were still undergoing division, expressed by

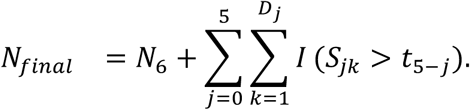

### Microglia Morphology

For microglial morphology analysis, 2-4 microglia were selected per animal in an imaging session. For each microglia selected a z-projection was created using FIJI. All microglia processes were manually traced, thresholded to generate a binarized outline of the process arbor, filtered to remove artifacts and analyzed with an automated Sholl Analysis plugin (provided by the Anirvan Ghosh laboratory, University of California, San Diego). The maximum number of intersections and the area under the curve (AUC) of the Sholl profile was analyzed to determine microglia arbor complexity.

### Microglia Motility and Surveillance

For motility analysis, XYZT images consisting of 40-μm-deep z-stacks were collected every 5 minutes, 12 times for a total of 60 minutes. Single-image 10 μm Z-projections were created for each time point, and lateral motion artifact was corrected using the StackReg and TurboReg functions (http://bigwww.epfl.ch/thevenaz/stackreg/). After thresholding and binarizing of the maximum intensity projections for all time points together, overlays of consecutive time points (0-5 minutes, 5-10 minutes, etc.) were made so that white pixels represented stability. A custom Matlab algorithm was used to compare pixels across individual time points and across consecutive time points to generate a motility index (defined as the sum of all changed pixels divided by the unchanged pixels). Additional indices were generated including stability, as the proportion of extension pixels (green) in one overlay that became stable (white) in the subsequent overlay divided by the total extension (green) pixels in the first overlay. Conversely, an instability index was calculated as the proportion of stable (white) pixels in one overlay that became retracted (magenta) in the subsequent overlay divided by the total stable (white) pixels in the first overlay.

For the surveillance ratio, we z projected all 12 time points and compiled them into a stack. We then aligned the stacks and modulated the brightness/contrast to ensure that all processes are visible and background is minimal. The z-projected files were then thresholded. The thresholding parameters were chosen to capture most of the processes while minimizing background pixilation. The thresholded time points were used to calculate the area surveyed (surveillance ratio) by microglia during the 1-h imaging session. This was done by calculating the total number of pixels representing microglia divided by the total pixels in the field of view. The normalized surveillance was calculated as the number of white pixels present in the 1^st^ time point divided by the total number of white pixels.

### Microglia Soma Area Quantification

2-10 microglia were selected at random from high magnification images. Single-image 10 μm Z-projections were created for each time point, and lateral motion artifact was corrected using the StackReg and TurboReg functions (http://bigwww.epfl.ch/thevenaz/stackreg/). The polygon tool was used to outline the soma of microglia and the area was calculated in microns (μm).

### Laser Ablation

Laser ablation injuries were created by running a point scan for 8s at 780nm using ∼75mW at the sample. The microglia injury response was imaged by collecting z-stacks of 50-90 μm every 5 minutes. For analysis, Z-projections were all comprised of 10 μm of the stack, encompassing the approximate center of the ablation core. The file was converted to an AVI and subjected to analysis by a custom MATLAB script designed to calculate the movement of microglia processes towards the ablation core. Briefly, for each pixel at each time point the script generates a vector which estimates the magnitude and direction of motion of the pixel utilizing the Farneback method for estimating optic flow. For analysis, we only included vectors larger than 5 pixels of motion which were directed towards the ablation core to minimize noise. The magnitude of all the vectors at each time point was summed and normalized to the total number of pixels in the image. For the convergence analysis, the number of pixels that enter the core of the focal injury are summed. We quantified the area under the curve, and the maximum value of the normalized magnitude over the 1-hour session.

### Histology

Whole brains were perfused with 0.1M PBS and fixed overnight with paraformaldehyde (4%). The tissue was cut on a freezing microtome (Microm; Global Medical Instrumentation) at 50 µm. For immunohistochemistry, sections were rinsed and endogenous peroxidase activity and nonspecific binding were blocked with a 10% BSA solution. Sections were then incubated in primary antibody solution to detect microglia (24h, 4 °C, anti-Iba-1, 1:1,000, Wako 019-19741) followed by secondary antibody solution (4 h, RT, AlexaFluor 488, 1:500, Invitrogen), mounted and coverslipped. To determine microglial depletion and repopulation, primary visual cortex sections were imaged on a Zeiss LSM 510 confocal microscope (Carl Zeiss). For each section, a 10-mm *z* stack in the center of the tissue was collected with a *z* step of 1 µm at x40 magnification. Analysis was performed offline in ImageJ. *Z* stacks were smoothed and compressed into a single *z* projection. Microglial cell bodies were marked in ImageJ using the paintbrush tool. Results from 4-5 sections per animal were averaged. Density was calculated as the number of microglia per area in visual cortex for PLX depleted and control groups.

For quantification of astrocytes, tissue was processed for histology as described above. Sections were incubated in a primary antibody solution (overnight, 4 °C, Anti-Glial Fibrillary Acidic Protein (GFAP), 1:500, Sigma G3893, clone G-A-5) followed by a secondary antibody solution (4 h, RT, AlexaFluor 594, 1:500, Invitrogen), mounted and coverslipped. To determine astrocyte coverage, somatosensory cortex sections were imaged on a Zeiss LSM 510 confocal microscope (Carl Zeiss). For each section, a 10-mm *z* stack in the center of the tissue was collected with a *z* step of 1 µm at x40 magnification. Analysis was performed offline in ImageJ. The images were thresholded and the total number of white pixels were then measured. Results from 3-4 sections per animal were averaged.

### Statistics

Statistical comparisons were made between animal and treatment groups using Prism V1 software (GraphPad, San Diego, CA). No statistical tests were used to determine sample sizes but our samples sizes are similar to those reported in the field. All *n* represents individual animals. For analysis were a number of microglia were selected and analyzed per animal, all microglia were averaged to generate a single value per animal. Animals were excluded from analysis if an imaging session could not be completed in full; otherwise, all completed imaging sessions collected were used. Not all animals were imaged at all imaging time points. All values reported are the mean ± s.e.m. For all analyses, *α* = 0.05. Two-tailed unpaired or paired *t*-tests and one-way or two-way ANOVA with or without repeated measures (ANOVA) with Tukey post hoc comparisons were used to compare cohorts where appropriate. The data met the assumptions of normality and equal variances as tested by Prism VI as part of the statistical analyses.

## Data Availability

The data that support the findings of this study are available from the corresponding author upon reasonable request.

## Code Availability

All MATLAB code is available at https://github.com/majewska-lab.

## Acknowledgments

We thank all the members of the Majewska lab for critical discussion and feedback on the manuscripts. We also thank the Olschowka and O’Banion lab for their valuable discussion and feedback. We thank Plexxikon for providing the PLX5622 drug for the depletion studies. This work is supported by grants from the National Institute of Health (NIH): F99 NS108486-02 (to M.S.M), R01 EY019277, RO1 NS114480 and R21 NS099973 (A.K.M), and NSF 1557971 (A.K.M.).

## Author contributions

M.S.M and A.K.M designed the study and wrote the manuscript with help from all authors. M.S.M performed the awake chronic *in vivo* two photon imaging experiments, motility and laser ablation experiments; J.A performed data analysis; and M.S.M wrote Matlab code to analyze the data; Z.B, M.S.M and M.M created, ran the Mathematical simulation and helped with statistical analysis.

