## Supplemental Figures for "*In vivo* imaging of the kinetics of microglial self-renewal and maturation in the adult visual cortex"

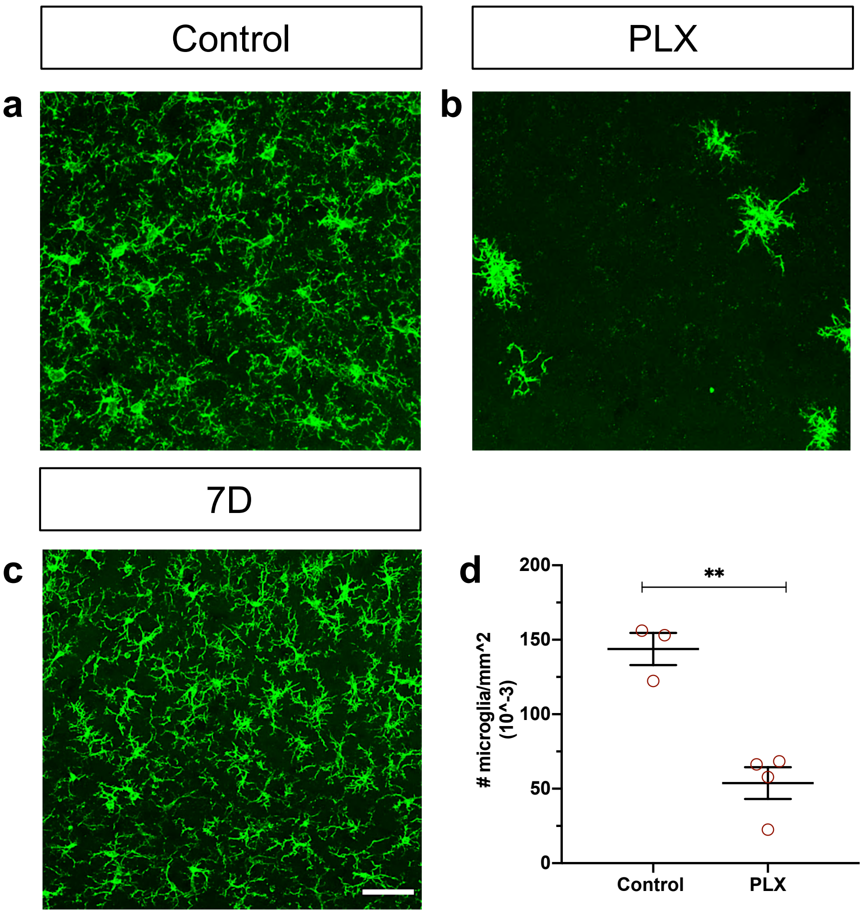


**Supplementary Figures 1: CSF1R inhibitor consistently eliminates microglia from the adult brain.** (a-c) Representative maximum intensity projections of confocal images of microglia from the primary visual cortex (V1) in fixed sections of control (a), 7 days PLX (b) and 7 days repopulated (c) mice. (d) Microglial numbers significantly decreased with PLX. n=3-4. with3-4 slices per animal. T-test, **p<0.01. Graphs show mean ± s.e.m; ns. Points represent individual animals. Scale bar, 50μm.

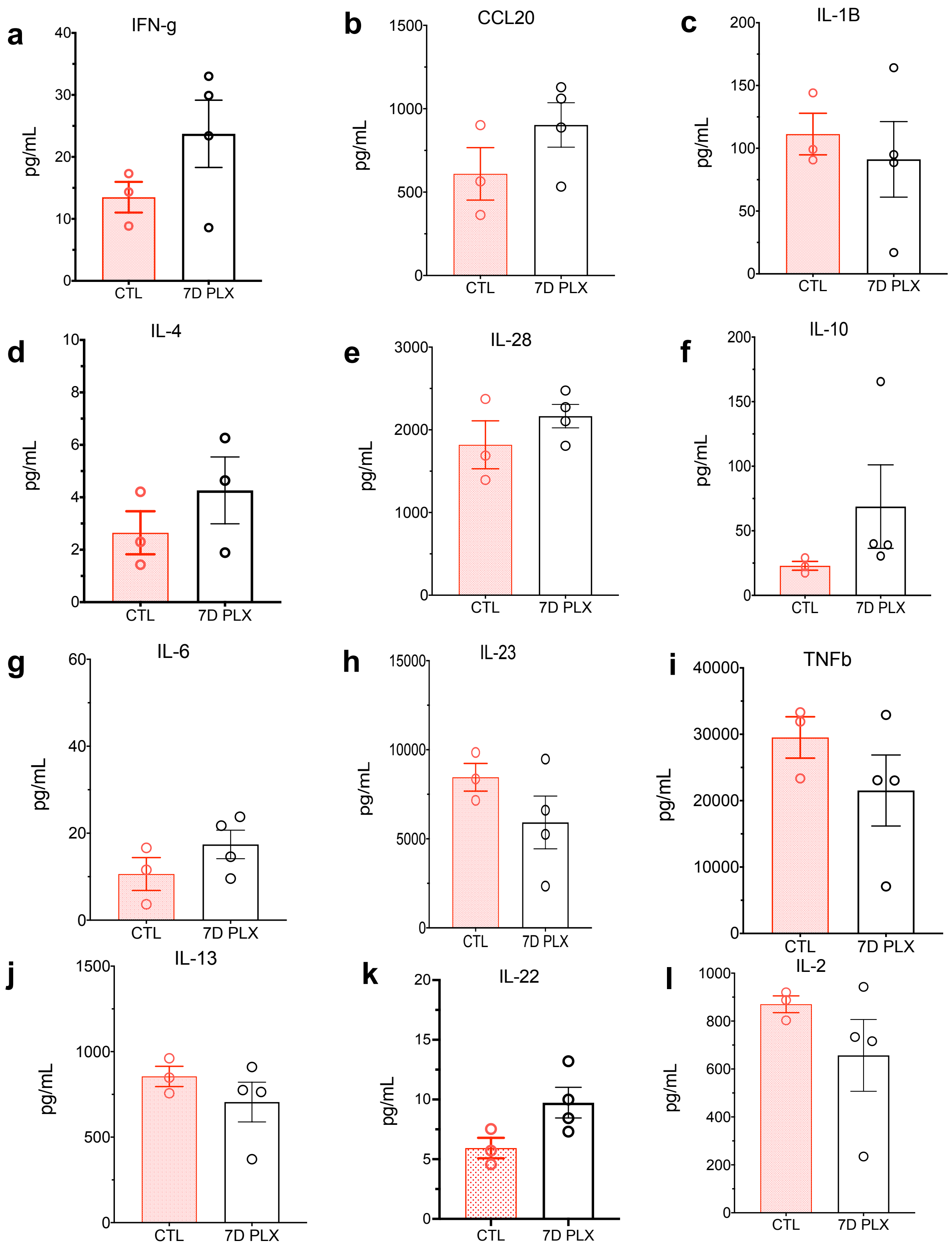

a
IFN-g
pg/mL
0
10
20
30
40
CTL
7D PLX
b
CCL20
pg/mL
0
500
1000
1500
CTL
7D PLX
c
IL-1B
pg/mL
0
50
100
150
200
CTL
7D PLX
d
IL-4
pg/mL
0
2
4
6
8
10
CTL
7D PLX
e
IL-28
pg/mL
0
1000
2000
3000
CTL
7D PLX
f
IL-10
pg/mL
0
50
100
150
200
CTL
7D PLX
g
IL-6
pg/mL
0
20
40
60
CTL
7D PLX
h
IL-23
pg/mL
0
5000
10000
15000
CTL
7D PLX
i
TNFb
pg/mL
0
10000
20000
30000
40000
CTL
7D PLX
j
IL-13
pg/mL
0
500
1000
1500
CTL
7D PLX
k
IL-22
pg/mL
0
5
10
15
20
CTL
7D PLX
l
IL-2
pg/mL
0
200
400
600
800
1000
CTL
7D PLX

**Supplementary Figures 2: No Change in inflammatory milieu following microglial depletion.** (a-l) A combination of anti-inflammatory and pro-inflammatory cytokines including IFN-g, CCL20, IL-1b, IL-4, IL-28, IL-10, IL-23, IL-6, TFN-b, IL-13, IL-22 and IL-2 were tested using\a MILIPLEX MAP mouse TH17 Magnetic Band Panel-Immunology Multiplex Assay. Each experiment was assayed in triplicate. Data are means +/- SEM of n=3-4 animals in each group (control and 7 days after PLX treatment). t-test, ns.

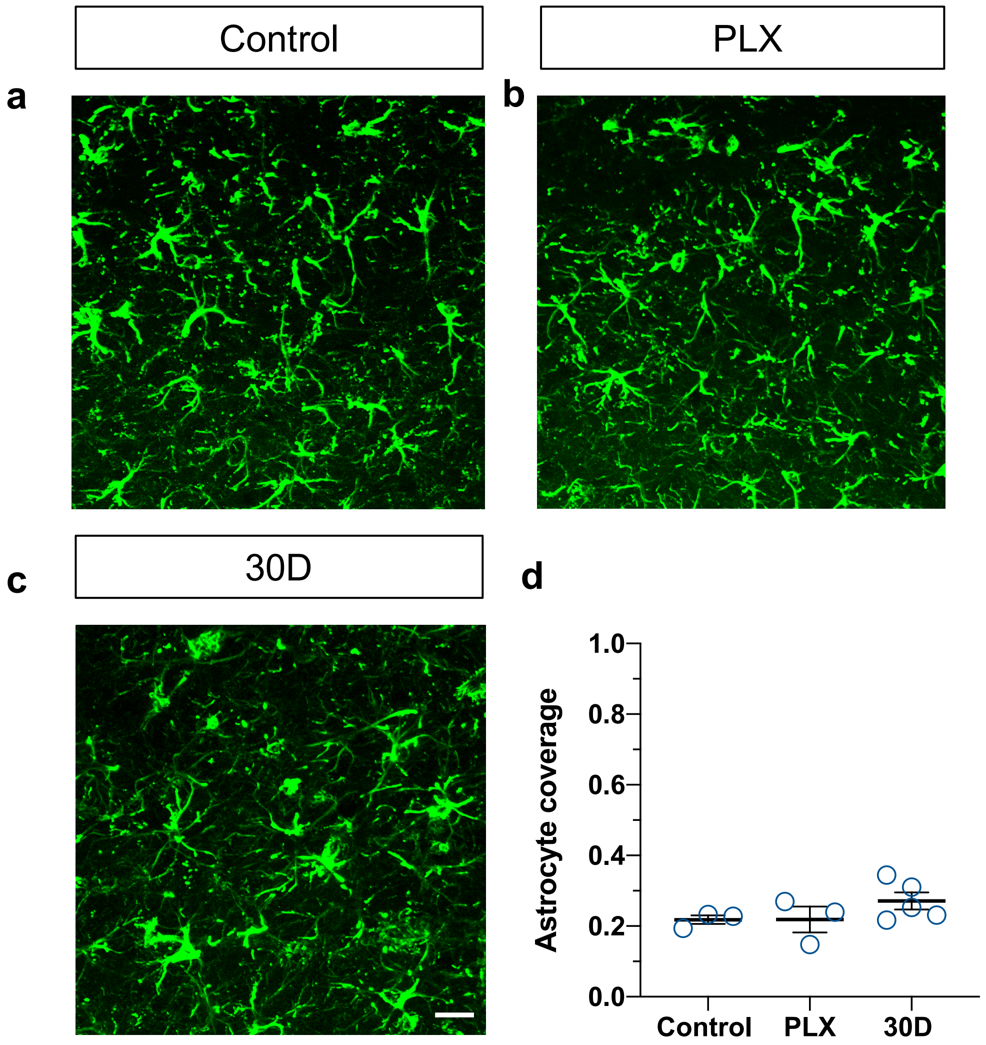


**Supplementary Figures 3: Astrocyte coverage of V1 is unchanged following microglial depletion.** Representative confocal images of astrocytes immunoreacted for GFAP in fixed brain sections from mice in control conditions (a) after PLX treatment (b) and after 30 days of repopulation (c). Qualitatively astrocyte morphology and expression of GFAP did not change following microglial depletion. (d) The proportion of V1 area covered by GFAP-positive astrocytes was unchanged with PLX treatment. n=3-5, ns, one-way ANOVA, Dunnett post hoc test. Scale bar, 20μm.

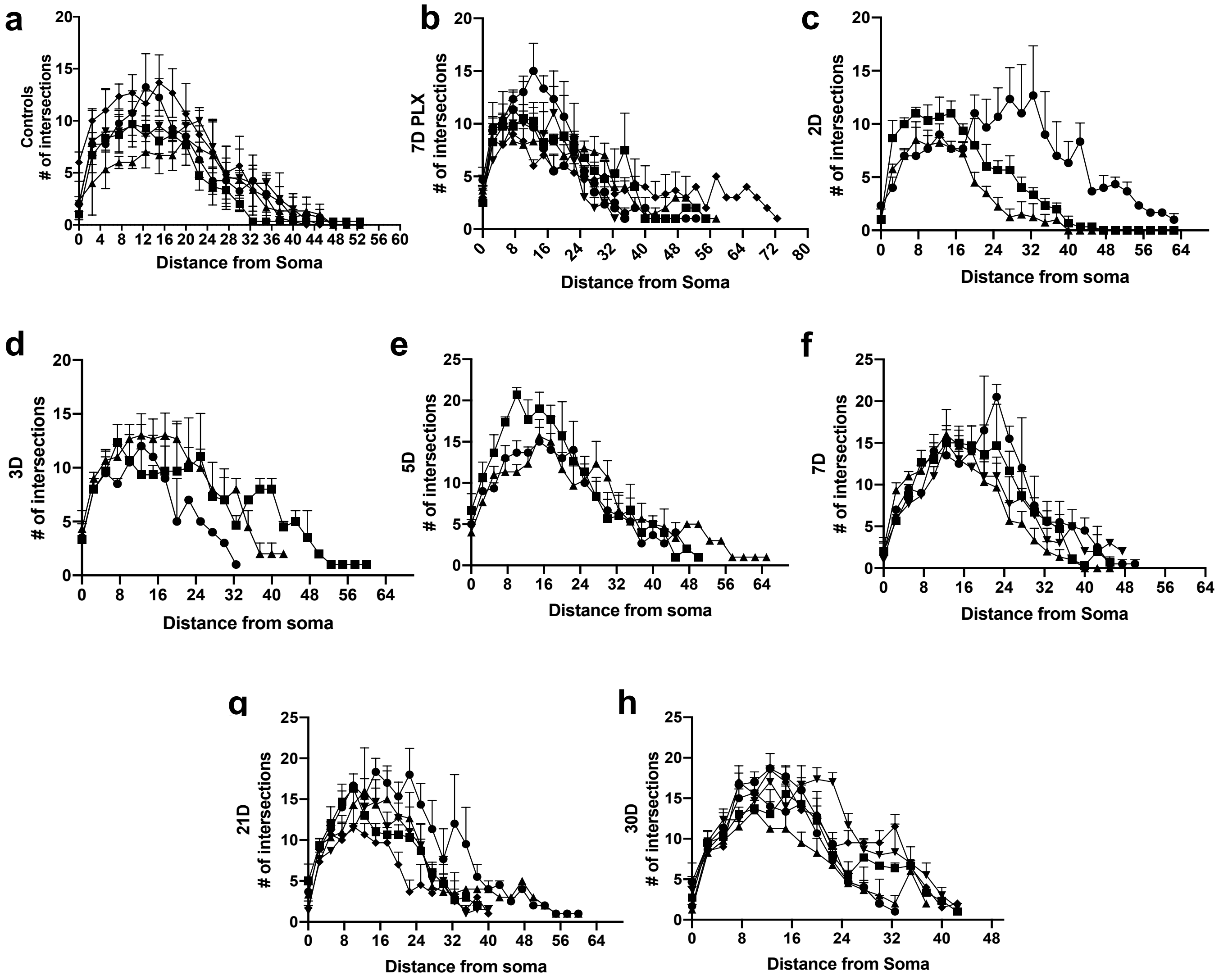


**Supplementary Figures 4: Microglial morphology over the course of repopulation.** (a-h) Average Sholl profiles from individual animals for each imaging time point (control day, 7D PLX, 2 days, 3 days, 5 days, 7 days, 21 days and 30 days repopulation; n=3-6 mice per group, 2-5 microglia per mouse, each line represents a different animal).

### Supplementary Videos:

**Supplementary Video 1: Example *in vivo* two photon imaging in awake adult mice.** Two-photon *in vivo* images were obtained in $CX_3Cr1^{GFP/+}$ awake mice following 3 consecutive days of training and habituation to the setup with a 1.5x zoom. This video shows a 100-slice z-stack starting at the pial surface and going deeper into the brain in a control awake animal. Microglia are evenly spaced and maintain this organization over many imaging sessions under control conditions. (Microglia are green). Scale bar, 50µm.

**Supplementary Video 2: Example *in vivo* two photon imaging in awake adult mice during depletion and repopulation.** Two-photon *in vivo* images were obtained in $CX_3Cr1^{GFP/+}$ awake mice following 3 consecutive days of training and habituation to the setup with a 1.5x zoom. This video shows a 100-slice z-stack in the same region during depletion (2D, 4D, 6D and 7D PLX) and with repopulation (D1-D7, D14, D21 and D30). Scale bar, 50µm.

**Supplementary Video 3: Example *in vivo* laser ablation video during depletion and repopulation.** Imaging was done in $CX_3Cr1^{GFP/+}$ awake with a 1.5x zoom. Here, we compared a laser ablation during control condition, depletion (PLX) and repopulation (D7) and D30). Microglia responded rapidly during these two conditions. 48 slices total, 12 slices per group. Scale bar, 20µm.
